# The dynamics of human cognition: increasing global integration coupled with decreasing segregation found using iEEG

**DOI:** 10.1101/089326

**Authors:** Josephine Cruzat, Gustavo Deco, Adrià Tauste, Alessandro Principe, Albert Costa, Morten L. Kringelbach, Rodrigo Rocamora

## Abstract

Cognitive processing requires the ability to flexibly integrate and process information across large brain networks. More information is needed on how brain networks dynamically reorganize to allow such broad communication across many different brain regions in order to integrate the necessary information. Here, we use intracranial EEG to record neural activity from 12 epileptic patients while they perform three cognitive tasks in order to study how the functional connectivity changes to facilitate communication across the underlying network spanning many different brain regions. At the topological level, this facilitation is characterized by measures of integration and segregation. Across all patients, we found significant increases in integration and decreases in segregation during cognitive processing, especially in the gamma band (50-90 Hz). Accordingly, we also found significantly higher level of global synchronization and functional connectivity during the execution of the cognitive task, again particularly in the gamma band. More importantly, we demonstrate here for the first time that the modulations at the level of functional connectivity facilitating communication across the network were not caused by changes in the level of the underlying oscillations but caused by a rearrangement of the mutual synchronisation between the different nodes as proposed by the “Communication Through Coherence” Theory.

## 1. Introduction

Intracranial electroencephalography (iEEG) recordings from the human brain provide a unique opportunity to study cognitive functions measuring neural activity at the mesoscopic level. Beyond the high temporal resolution intrinsic to EEG measurements, this technique allows also higher levels of spatial resolution and enhanced signal-to-noise ratio [1, 2]. These advantages have led scientists to use the technique [3] to study several cognitive processes such as attention [4], visual perception [5, 6], language [7–9], memory [10–12], decision making [13], emotion [14, 15] and consciousness [16]. However, most of this research has attempted to assign functions to specific local brain areas by correlating task-performance with measurements of neural activity - but see also recent coherence network studies [17–19]. Furthermore, most studies using iEEG have focused mainly on single-electrodes analysis, using predominantly event-related potentials [3] or spectral analysis [20].

In contrast to this regional view of brain function, growing evidence reveals that human cognition relies on the flexible integration of information widely distributed across different brain regions [21, 22]. Many studies have investigated the brain networks’ properties underlying cognitive processing using EEG [23, 24], MEG [25–27] and fMRI [28–32]. These studies concur in that cognitive processing appears to increase the global integration of information across neural networks, while at the same time leads to a decrease in the modularity of those networks [25, 30, 31, 33–36]. These findings, have not been validated in iEEG studies yet. More importantly, there is still a lack of knowledge about the underlying mechanisms causing the cognitively driven modulation of the level of integration and segregation across the network.

The “Communication Through Coherence” (CTC) theory states that the synchronisation between different neuronal populations could modulate the communication and information processing between them [37]. Indeed, two populations of neurons may communicate most effectively, when they are coherent, i.e. when their excitability level is coordinated in time. Indeed, the CTC theory suggests that effective connections in a network can be shaped through phase relations, more specifically through gamma- and beta-band (30–90 Hz) synchronisation, as reported experimentally [38–40]. Thus, oscillations are proposed to dynamically shape the computational role of different neuronal groups linked through static structural connectivity. Several empirical studies support task-induced increases in synchronisation at the level of individual regions during selective attention [41], working memory [42], and motor control [43]. Moreover, such task-induced changes in synchronisation have been observed between distant cortical regions during working memory [44], long-term memory encoding [45], visual attention [46], and sensorimotor integration [47]. The main aim of this paper is to link the CTC theory with modulations of the level of integration across different brain areas during human cognition. Here, we explore how cognitive processing modulates the level of integration and segregation of information in human brain networks. More specifically, we recorded iEEG data from depth electrodes stereotactically implanted for pre-surgical diagnosis in 12 epileptic patients performing three different cognitive tasks. The iEEG electrodes used a stereoelectroencephalography (SEEG) implantation methodology and covered broad regions of the brain including cortical as well as subcortical regions, so that we were able to assess the global changes at the level of a broad extended network. This allowed us to investigate the network properties using the operationally defined concepts of segregation and integration [22] as global network measures of brain function. Furthermore, we also assessed how these concepts relate to synchronization rather than to the oscillations level (amplitude). Finally, we would like to emphasize that our claims about the modulation of integration/segregation and the validity of CTC theory in our research are global, and it is because our statements are independent of the electrodes placement (i.e. node location, which was different for each patient) and type of cognitive processing (three different cognitive tasks were used).

## 2. Material and Methods

### 2.1 Ethics Statement

The Clinical Research Ethical Committee of the Municipal Institute of Health Care (CEIC-IMAS) approved this study. Following the Declaration of Helsinki, patients were informed about the procedure and they gave their written consent before the experiment.

### 2.2 Participants

Twelve participants (3 women; all right-handed; mean age 36.4±10.1 years-old), evaluated for presurgical diagnosis in the Epilepsy Monitoring Unit of the Hospital del Mar (Barcelona, Spain), participated in the study. All patients were stereotactically implanted with depth electrodes for invasive presurgical diagnosis using a stereotactic ROSA robotic device (Medtech, France). The location of the electrodes was established only for clinical reasons using a SEEG approach. The implantation schemas were similar between all patients given that they were all under investigation for temporal lobe epilepsy. The number of electrodes used varied among 8 to 16 for patient with 5 to 15 contacts each (diameter: 0.8 mm; contacts 2 mm long, 1.5 mm apart) (DixiMédical, France). All patients underwent an extensive neuropsychological evaluation, and had normal or corrected-to-normal vision. They were within the normal range of education having completed from primary to high academic level. Table 1 summarizes personal data, pathological information and overview of implanted electrodes for each patient. Since the study aims to study the network dynamics supporting cognitive processes under normal circumstances, patients were assessed in absence of pharmacological treatment.

**Table 1.** Demographic and clinical characteristics of each patient.

| Patient | Gender | Age (years) | Epileptogenic zone laterality | Seizure onset zone | Implanted Regions | N° of electrodes implanted | N° unipolar channels | N° of bipolar channels for analysis |
| --- | --- | --- | --- | --- | --- | --- | --- | --- |
| A | Male | 38 | Left | Mesial temporal and amygdala | L (F-T-I-P) | 8 | 85 | 76 |
| B | Male | 21 | Left | Anterior temporal | L (F-T-I-P-O) & R (F-T) | 15 | 125 | 95 |
| C | Male | 44 | Right | Anterior temporal | L (T) & R (F-T-P) | 12 | 125 | 104 |
| D | Male | 44 | Right | Temporo-parietal | L (T) & R (F-T-I-P) | 16 | 127 | 101 |
| E | Female | 43 | Left | Anterior medial temporal | L (F-T-I-P-O) | 11 | 127 | 112 |
| F | Male | 25 | Right | Temporo-parieto occipital | R (F-T-I-P-O) | 14 | 127 | 105 |
| G | Male | 23 | Left | Temporal | L (F-T-I-P) | 13 | 125 | 102 |
| H | Female | 43 | Left | Posterior Temporal | L (F-T-I-P) | 14 | 125 | 103 |
| I | Male | 44 | Left | Mesial Temporal | L (F-T-I-P) | 12 | 123 | 105 |
| J | Female | 46 | Left | Anterior medial temporal | L (F-T-I-P-O) | 11 | 124 | 107 |
| K | Male | 43 | Left | Mesial Temporal | L (F-T-P) | 9 | 103 | 94 |
| L | Male | 23 | Right | Anterior temporal | R (F-T-P) | 15 | 125 | 96 |

R: Right; L: Left; F: Frontal; T: Temporal; P: Parietal; O: Occipital; I: Insula

### 2.3 Cognitive Tasks

#### 2.3.1 Picture-Naming Task

In the picture-naming task, participants were asked to name 228 pictures presented in three different blocks (N=228). Pictures were black & white line drawings of familiar objects from a wide range of semantic categories selected from the Snodgrass and Vanderwart (1980) set. Each picture appeared once centrally and sequentially on the computer screen in a pseudo-random order for 2000 ms followed by a fixation cross for 1000 ms (see Figure 1). Participants were instructed to overtly name every item as fast and accurately as possible in Spanish.

**Figure 1.**
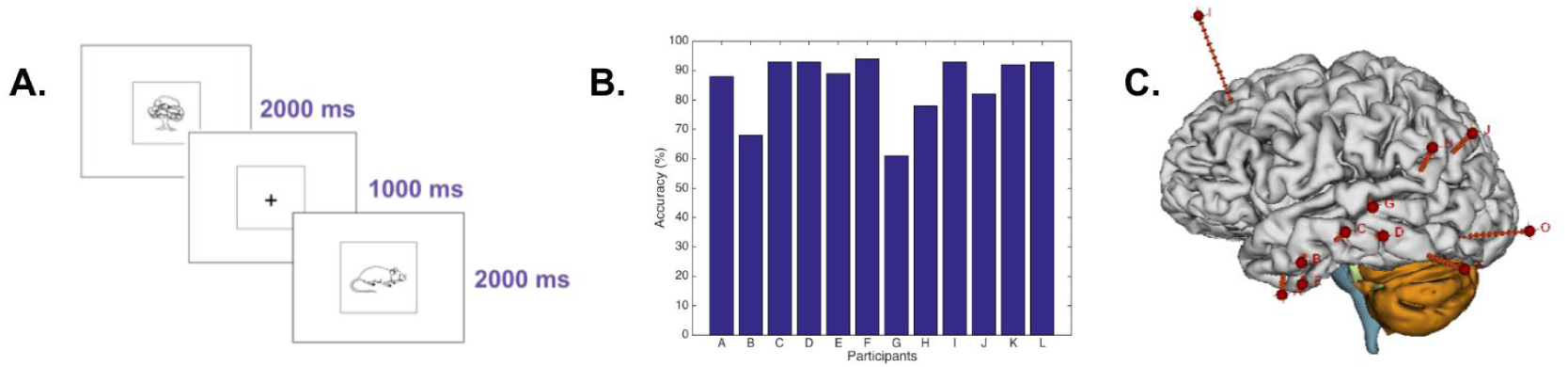
Paradigm, behavioural performance and implantation scheme. A) Schematic example of the experimental paradigm of the picture-naming task. After an interval of 1000 ms for preparation, each target picture was presented and remained on the screen for 2000 ms. B) Picture naming accuracy for each patient. On average, patients achieved 85.3±11.0% accuracy on test items (N=228). C) Example of the intracerebral implantation scheme for iEEG recordings in patient E. Eleven electrodes were implanted in the left hemisphere.

#### 2.3.2 Size-Judgement Task

Participants were instructed to indicate via button press if the presented Spanish word represented an object larger than one-foot box in any dimension. Word stimuli were 68 items from two different categories: animals and man-made objects. Half of the words in each category represented objects larger than one foot. Words were auditory-presented through the computer speakers. Each word was presented sequentially in a pseudo-random order followed by an inter-trial lapse of 3500 ms.

#### 2.3.3 Lexical-Decision Task

Participants were instructed to indicate whether the letter string presented in each trial written on a computer screen was a real word (e.g. run, table) or a pseudo word (e.g. lun, tible). We included four types of real words: motion verbs (e.g. run), static verbs (e.g. think), concrete nouns (e.g. table) and abstract nouns (e.g. theory). Each word was presented once centrally and sequentially on the computer screen in a pseudo-random order for 2000 ms followed by a fixation cross for 1000 ms.

The three tasks had different presentation modality of the stimuli, which allowed us to compare the pattern of intracranial EEG response elicited when retrieving conceptual knowledge from a lexical form (size-judgment task) with the activity elicited when retrieving the lexical form of an object depicting a concept (picture naming task) and with the one elicited when retrieving the concept through a written lexical form (lexical-decision task).

The three tasks were presented using the software Sketchbook Processing 2.2.1 (Programming Software, 2001 https://processing.org/) on a laptop computer at an approximate size of 5 degrees of visual angle. For the size-judgement and lexical-decision task, responses were given through a joystick connected to the computer. The accuracy of the responses for the picture-naming task was scored manually by the experimenter and for the other two tasks; the software collected response latencies and accuracy. An electronic processor “Arduino, UNO” was used to connect and synchronize both hardware; the XLTEK system with the computer (MacBook Pro). The application interfaced with an Arduino board that in turn was connected to the EEG amplifier, and at each trial a signal was sent through the Arduino to the EEG.

### 2.4 iEEG Data Acquisition and Pre-processing

Neurophysiological responses were registered by the iEEG system from deep multichannel electrodes (DIXI Microtechniques, Besançon, France). On average, each patient had 13±2 electrodes implanted (range 8-16) with a total of 120 ± 13 recording contacts (range 85-127). The data were acquired continuously by the Neuroworks XLTEK system (version 6.3.0, build 636) at 32 kHz with a headbox of 128 channels recorded at a sampling frequency of 500 Hz per channel. For our analysis, we considered only the channels placed in both cortex and subcortical structures. Channels placed in white matter or misplaced out of the cortex or subcortical structures were disregarded.

A bipolar montage was constituted offline to increase spatial resolution by removing any confounds from the common reference signal [2, 59]. Bipolar signals were derived by differentiating neighbouring electrode pairs of recorded and not rejected consecutive channels within the same electrode array [3, 15, 16, 60]. The continuous iEEG data was first high-pass filtered at 1 Hz and low-pass filtered at 150 Hz. To remove common line contamination an extra notch filter was applied at 50 and 100 Hz. In order to have specific spectral information, we analysed the spatio-temporal correlations of the Band Limited Power (BLP) at a given carrier frequency. This is a standard and successful approach introduced in the context of MEG analysis [61]. For that, the analysis at a given carrier frequency f_carrier_ (we consider here f_carrier_ = 1 to 130 Hz in steps of 4 Hz) requires first that the iEEG signals are band-pass filtered within the narrow band [f_carrier_-2, f_carrier_+2 Hz] (we used the second order Butterworth filter) and computed the corresponding envelope of each narrowband signal using the Hilbert transform [62, 63]. The Hilbert transform yields the associated analytical signals. The analytic signal represents a narrowband signal, s(t), in the time domain as a rotating vector with an instantaneous phase, φ(t), and an instantaneous amplitude, A(t), i.e., s(t)=A(t)cos(φ(t)). The phase and the amplitude (envelope of that carrier frequency) are given by the argument and the modulus, respectively, of the complex signal z(t), given by z(t)=s(t)+i.H [s(t)], where i is the imaginary unit and H[s(t)] is the Hilbert transform of s(t). We further consider only the slow components of the envelope A(t) by filtering the amplitudes again below 12 Hz [64]. Finally, the slow component of the envelope of each brain node, which corresponds to each bipolar channel, at a given carrier frequency was used to calculate the envelope FC.

In addition, we ran extra simulations (Figure 9) with a monopolar montage in order to test another possible definition of integration and to show the robustness of the results (independent of the montage). In this case, the brain nodes correspond to each single monopolar channel.

### 2.5 Data Analysis

#### 2.5.1 Envelope Functional Connectivity (FC)

For the three tasks, the data was segmented into two windows around stimulus presentation: the first one spanning from -500 ms to 0 (pre-stimulus window) from the stimulus presentation, and the second one spanning from 0 to 500 ms (post-stimulus windows) from the stimulus presentation. We defined an Envelope FC matrix of the continuous bipolar iEEG data for pre- and post-stimulus windows as a matrix of Pearson’s correlations of the corresponding amplitude envelopes, i.e. the slow components of the BLP of iEEG signals at a given carrier frequency between two brain areas over the whole time window for a given window (pre-stimulus and post-stimulus). Thus, the mean FC that we plot in the Results section is specific for each narrow band frequency window.

#### 2.5.2 Phase-Lock Matrix

For each time point we calculated the phase lock matrix describing the global state of synchronization across all network nodes using the monopolar montage. The elements of the phase-lock matrix are given by:

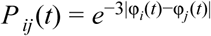

where φ_*i*_(*t*) is the extracted phase of node *i* at time *t* (at a given carrier frequency using the Hilbert transform as specified above). One can use the phase-lock matrix at a specific single time point to calculate the integration as specified below but instead of using the envelope FC using this single-time-point phase-lock matrix. For other simulation, where we contrast the pre- and post-stimulus windows, we considered the averaged phase-lock matrix across the time elapsed in the respective windows.

#### 2.5.3 Integration

We used the measure of integration introduced by Deco et al. (2015), defined at the network level, to characterize the level of broadness of communication between regions across the whole brain. First, we filtered the data, and then we calculated the envelope FC as explained above for both pre- and post-stimulus windows. We define integration as the size of the largest connected component in the FC matrix. That is, the number of nodes of the largest connected graph in the binarized functional connectivity matrix obtained after thresholding. More specifically, for a given absolute threshold θ between 0 and 1 (scanning the whole range), the FC (using the criteria |FC_ij_|<θ, i.e a value of 0 and 1 otherwise) can be binarized and the largest subcomponent extracted as a measure of integration. Note, that we only took absolute values because we want to account for the correlation independent if it is negative or positive. In graph-theoretical terms, subcomponents are extracted from the undirected graph defined by the binarized matrix (which itself is considered as an adjacency matrix). More precisely, a subcomponent is a subgraph in which paths connect any two vertices to each other, and which connects to no additional vertices in the super-graph[22]. A vertex *u* is said to be connected to a vertex *v* in a graph *G* if there is a path in *G* from *u* to *v*. The concepts of subgraph and super-graph are defined as following: Let *H* be a graph with vertex set *V(H)* and edge set *E(H)*, and similarly let *G* be a graph with vertex set *V(G)* and edge set *E(G)*. Then, we say that *H* is a subgraph of *G* if *V(H)* ⊆ *V(G)* and *E(H)* ⊆ *E(G)*. In such a case, we also say that *G* is a super-graph of *H*. Finally, to get a measure that is independent of the threshold, this curve can be integrated in the range of thresholds between 0 and 1. This integration measure is normalized by the maximal number of connected brain areas (that is, all N areas) for each integration step and by the number of integration steps such that the maximal integration is normalized to 1.

#### 2.5.4 Segregation

As a complement of the integration, we used the modularity measure [56] as a measure of segregation. Following Rubinov and Sporns (2011), modularity is defined as a measure of the goodness with which a network is optimally partitioned into functional subgroups, i.e. a complete subdivision of the network into non-overlapping modules, and supported by densely connected network communities. We consider the modularity of our envelope FC matrix. This matrix contains positive and negative weights, namely the corresponding correlation between two nodes. Our measure of modularity is given by,

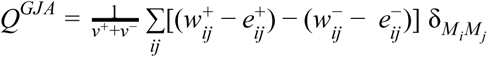

Where the total weight, v^±^ = Σ_ij_w_ij_^±^, is the sum of all positive or negative connection weights (counted twice for each connection), being w_ij_^+^ ∈ (0,1] the weighted connection between nodes i and j. The chance-expected within-module connection weights 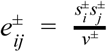, where the strength of node i, s_i_^±^ = Σ_j_w_ij_^±^, is the sum of positive or negative connection weights of i. The delta δ_MiMj_ = 1 when i and j are in the same module and δ_MiMj_ = 0 otherwise [65]. This definition is a generalisation of the standard measure of modularity for matrices with nonnegative weights, which is given by the average difference between present within-module connection weights w_ij_^+^ and chance-expected within-module connection weights e_ij_^+^. As mentioned above, here we consider both positively and negatively weighted connections (envelope FC matrix). The positively weighted connections represent correlated activation patterns and hence to reinforce the placement of positively connected pairs of nodes in the same module. On the other hand, the negatively weighted connections represent anti-correlated activation patterns and reinforcing the placement of negatively connected pairs of nodes in distinct modules. For a complete description see [66].

#### 2.5.5 Synchronization

We measure the global mean level of synchronization as the mean value of the Kuramoto order parameter across time. The Kuramoto order parameter is defined by following equation:

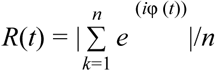

where φ_k_(t) is the instantaneous phase of each narrowband signal at node k at a given carrier frequency by using the Hilbert derived phases of the slow component of the Band Limited Power (BLP) signals. The Kuramoto order parameter measures the global level of synchronization of the n oscillating signals. Under complete independence, the *n* phases are uniformly distributed and thus *R* is nearly zero, whereas for R=1 all phases are equal (full synchronization).

#### 2.5.6 Permutation Tests

To examine the reliability of the post-stimulus window modulation, frequency-dependent statistical significance was assessed using a non-parametric test with 1000 random permutations. The test statistic was chosen to be the median across the differences between post and pre-stimulus measures. Following Winkler et al. (2014), we simulated the null hypothesis of no stimulus modulation at any post-stimulus windows by randomly permuting pre- and post-stimulus samples at each paired trial (group exchangeability hypothesis) and computing the test statistic. These surrogate values formed a reference distribution, against which we compared the original statistic value. The proportion of permutations in which surrogate values matched or exceeded the original statistic value determined the test p-value (P) to be compared with the significance level. We obtained a p-value for each frequency band and applied multiple comparisons corrections with the False Discovery Rate (FDR) level 0.05 using the Benjamini-Hochberg method.

## 3. Results

In the present study we exploited the high spatiotemporal resolution of intracranial electroencephalography (iEEG) recordings to investigate the reorganization of brain networks driven by cognitive tasks. This technique is usually employed in pharmacologically resistant epileptic patients [48, 49] who require brain mapping before surgery, and also provide a useful tool for basic research on cognitive neuroscience [3].

We recorded iEEG data in twelve patients while performing a picture-naming task (Figure 1A). This task was used to drive the modulation of the underlying brain networks related to the integration of task-related information. It is known that in the large majority of people language is supported by a widespread large-scale network distributed across frontal, temporal, parietal and occipital lobes in the dominant hemisphere [29, 50, 51]. During the task, patients had to overtly name each picture as fast as possible in Spanish (see Materials and Methods). The picture naming accuracy was high (85.3 ± 11.0%) (Figure 1B). The averaged response time was 1350 ms. All channels recordings from grey matter and subcortical structures were considered for the analysis. Although neural activity signal was recorded from all lobes in both hemispheres, most of the recordings were obtained from the left temporal lobe (see an example of implantation scheme in Figure 1C). After pre-processing the data, we analysed the Band Limited Power (BLP) at a given carrier frequency (*f*_*carrier*_) in order to have specific spectral information. We band-pass filtered the iEEG signals within the narrow band [*f*_*carrier*_ -2, *f*_*carrier*_ +2 Hz] and considered a range of *f*_*carrier*_ starting from 1 to 130 Hz in steps of 4 Hz. We chose a bandwidth of 4 Hz because it provides a good trade-off between phase estimation accuracy and number of testable comparisons. In order to compute the envelope Functional Connectivity (FC), we further computed the Hilbert transform (Figure 2). To study changes in the functional network topology during the execution of a cognitive task, we contrasted two situations, namely: a pre-stimulus window of 500 ms before the picture was presented: *“pre-stimulus window”*, and a post-stimulus window of 500 ms immediately after the picture was presented: *“post-stimulus window”.* In order to characterize the organization of the network under both situations, we used the integration and segregation measures of global brain function (see Materials and Methods) characterizing the level of communication across the different nodes of the brain network. We tested the statistical significance of the network measure differences between pre-stimulus and post-stimulus using a non-parametric method [52]. Specifically, we characterized the null hypothesis using constrained permutations that preserved the trial grouping of the data at both time windows (Materials and Methods).

**Figure 2.**
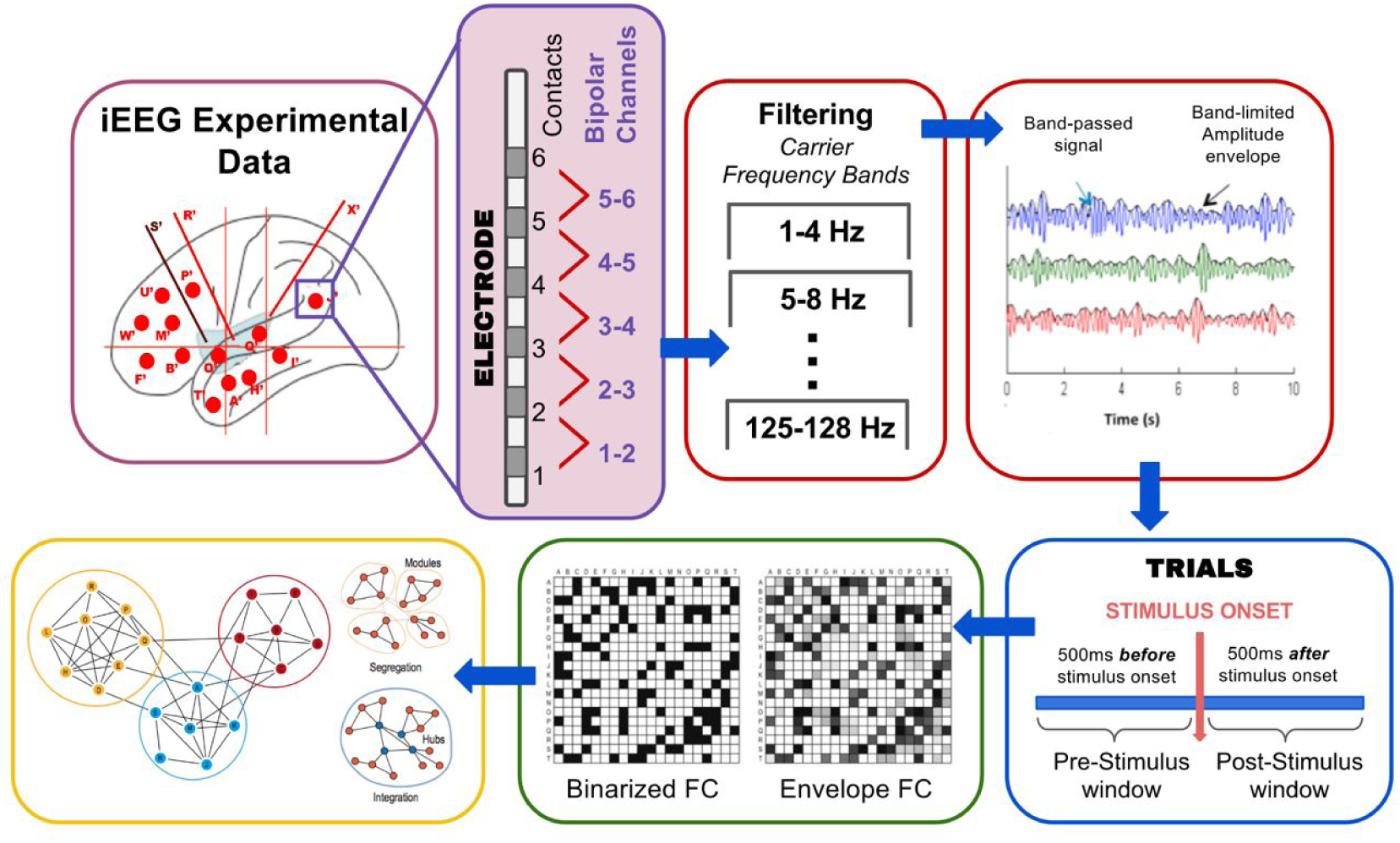
Data processing flow chart. The iEEG data is recorded from 85 up to 127 unipolar channels on each patient. The bipolar montage is constituted offline by subtracting the neural activity recorded by neighbouring contacts within the same electrode array. The data is first band-pass filtered at 1 to 150 Hz, and further band-pass filtered into narrow frequency bands [f_carrier_-2, f_carrier_+2 Hz] (we consider here carrier frequencies f_carrier_= 1 to 130 Hz in steps of 4 Hz). By the Hilbert transform the corresponding amplitude envelopes are computed to further compute the envelope FC matrix. The continuous data is segmented into windows of -500 to 0 ms (pre-stimulus window) and 0 to 500 ms (post-stimulus window), around stimulus presentation. In order to characterize the organization of the network under both windows, we used the integration and segregation measures of global brain function.

The main results come from the comparison of the integration and segregation measures during the pre- and post-stimulus windows. Here, we first focus on a single patient to illustrate the main results (Figure 3). Neural integration significantly increases during cognitive processing (Figure 3A) and complementary to this measure, we observed a consistent decrease in segregation as measured by the modularity (Figure 3B). In both cases, the greatest modulation appeared particularly in the gamma band, around 50 to 90 Hz (p<0.05, N=1000). Note that the concepts of integration and segregation are measures that by definition are calculated independently from each other but their results are highly correlated; when one increases the other consistently decrease.

**Figure 3.**
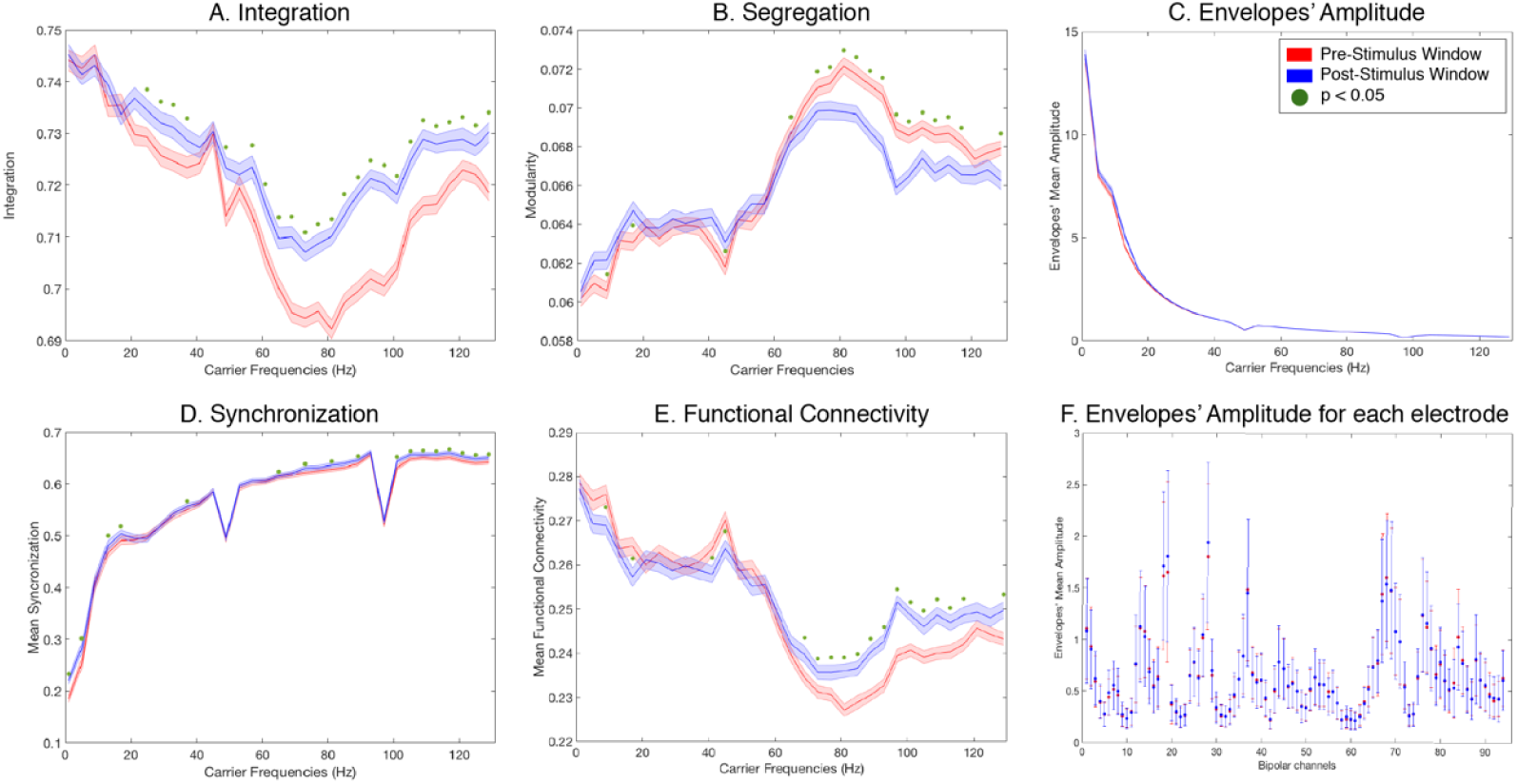
Results analysis patient K for the picture-naming task. Panel A shows a significant increase in the integration during cognitive processing. The greatest effect is observed in the gamma range (50 to 90 Hz). The red line corresponds to pre-stimulus window, the blue line corresponds to post-stimulus window, the error shaded regions reflect the standard deviation across trials and green dots indicates a statistical significance of p<0.05 (N=1000). Complementary to the integration, panel B shows a decrease of the modularity in the same frequency range. Panel C shows that there are no iEEG oscillations amplitude changes in any frequency induced by the stimulus (both curves are strongly overlapped). This result indicates that the increase of integration and decrease of modularity could not be explained by changes in the oscillations amplitude. Panel D shows an increase of mean synchronization over a broad range of frequencies that is more conspicuous in the gamma band range (50 to 90 Hz) for the post-stimulus window. Panel E shows that the functional connectivity behaves coherently with the other results, as it increases as a function of the stimulus presentation particularly in the gamma range (50 to 90 Hz). Panel F plots the amplitude of the oscillations envelope at 60 Hz for each bipolar channel and pre- and post-stimulus window. There are no noticeable modulations across single bipolar channels between pre- and post-stimulus window. Note that the sharp peaks at 50 and 100 Hz are due to the power-line noise created by the electrical power.

Having found an increase in the integration consistently associated with a decrease in the segregation, the question then arises what is behind the modulation of these measures driven by the task. We first examined whether these modulations could be explained by changes in the oscillations’ envelope amplitude. Importantly this was not the case; no modulation of oscillation amplitude at any frequency (Figure 3C) could explain either the integration increases or the modularity decrease. This was so both when looking at the mean amplitude of all electrodes together and when evaluating the mean amplitude of each bipolar channel separately at 60 Hz (which corresponds to the maximal modulation of the integration measure) (Figure 3F).

Further, we looked whether the level of global synchronization of the corresponding envelopes was associated with the modulation of the integration and segregation. We used the Kuramoto Order Parameter (see Materials and Methods) to measure the mean synchronization of cortical activity under both conditions. We observed that, in fact, there is an increase of mean synchronization over a broad range of frequencies that is greater in the gamma range (50 to 90 Hz) for the post-stimulus window (Figure 3D). Next, we computed the Envelope Functional Connectivity (see Materials and Methods) between channels to see whether it behaves coherently with the level of global synchronization. We calculated it through the instant amplitude envelopes of the given frequencies. The analysis revealed that the Envelopes’ Functional Connectivity is enhanced during task performance, in particular in the gamma range (50 to 90 Hz) (Figure 3E).

Together these results suggest that given that the relationship between integration and segregation cannot be accounted by an increase of the oscillations, the most likely explanation for these phenomena is a global increase of the connectivity, especially in the gamma band. This is consistent with the notion that communication between different brain networks is accomplished by means of the level of synchronization as posited by the *Communication Through Coherence Theory (CTC)* [38, 40].

Importantly, the significant integration and segregation modulations driven by stimulus presentation were consistently found in *every patient*. Figures 4 and 5 plot for each single patient across windows, the cognitive modulation of integration and segregation, respectively. As shown in the figures, despite the heterogeneity of the recording sites, all patients show consistently the same pattern of modulation. Moreover, the increase of integration was correlated with the decrease of segregation, as it was shown quantitatively by the Pearson correlation of -0.9846 (N=12) of the respective modulations at the carrier frequency of 60 Hz (maximal modulatory effect point). Interestingly, in most of the patients, there was a consistent decrease of the segregation in the gamma band but with a slight increase of the segregation in the sub-gamma regime. This is not observed for the integration measure. Figure 6 plots the oscillations’ envelope amplitude at 60 Hz for each bipolar channel and for each patient. In all cases, the amplitude of the oscillations envelope across electrodes does not differ between pre- and post-stimulus windows. Indeed, the majority of the electrodes (over 98.6%) do not show significant differences in amplitude modulation.

**Figure 4.**
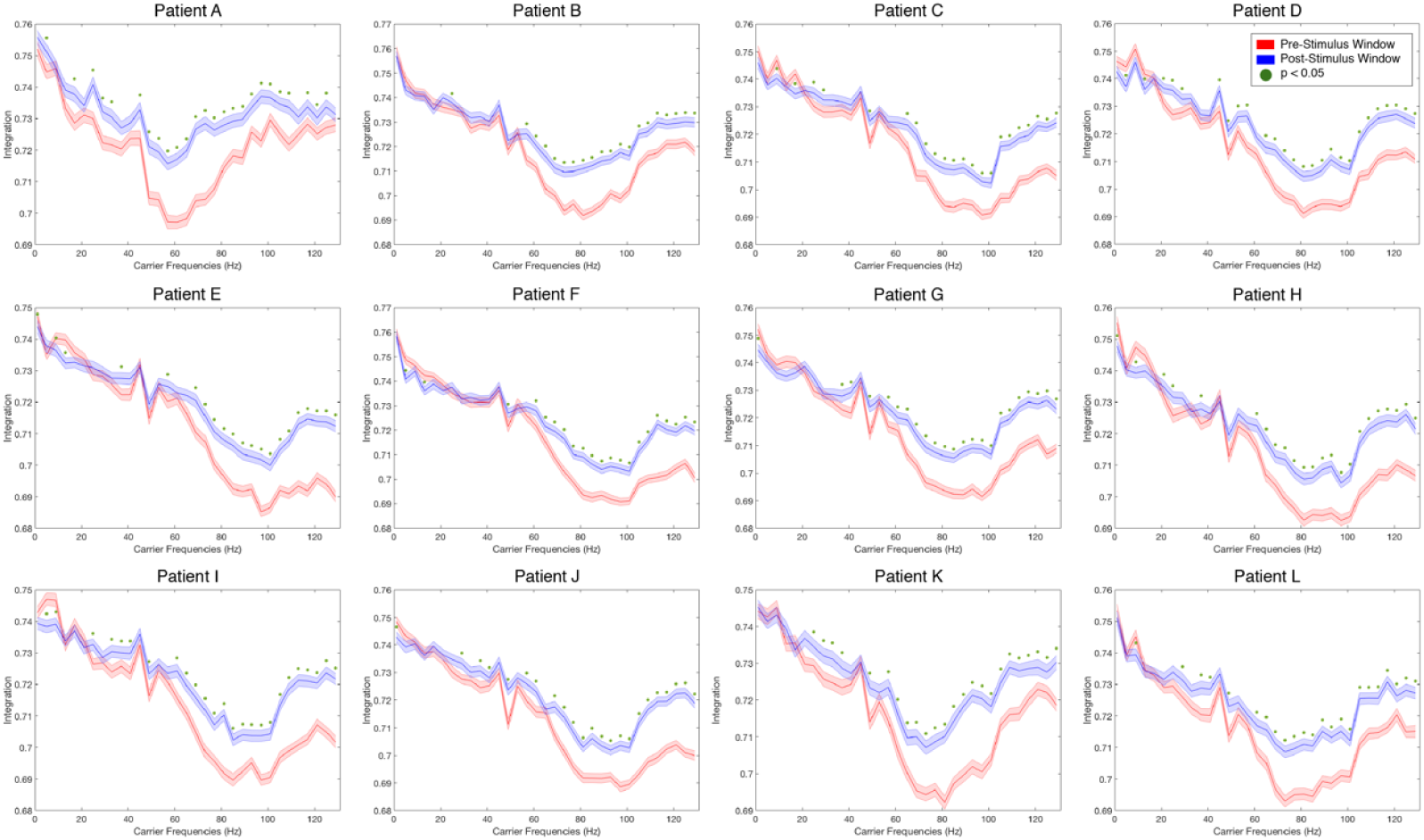
Integration measure results for each patient. The panels show the results for every single patient for the picture-naming task. As can be seen, despite the heterogeneity of the recording sites, all patients show a significant increase of the integration related to cognitive processing. For all patients, the greatest effect was found in the gamma range. The red line corresponds to pre-stimulus window, blue line corresponds to task condition, the error shaded regions reflect the standard deviation across trials and green dots indicates a statistical significance of p<0.05 (N=1000).

**Figure 5.**
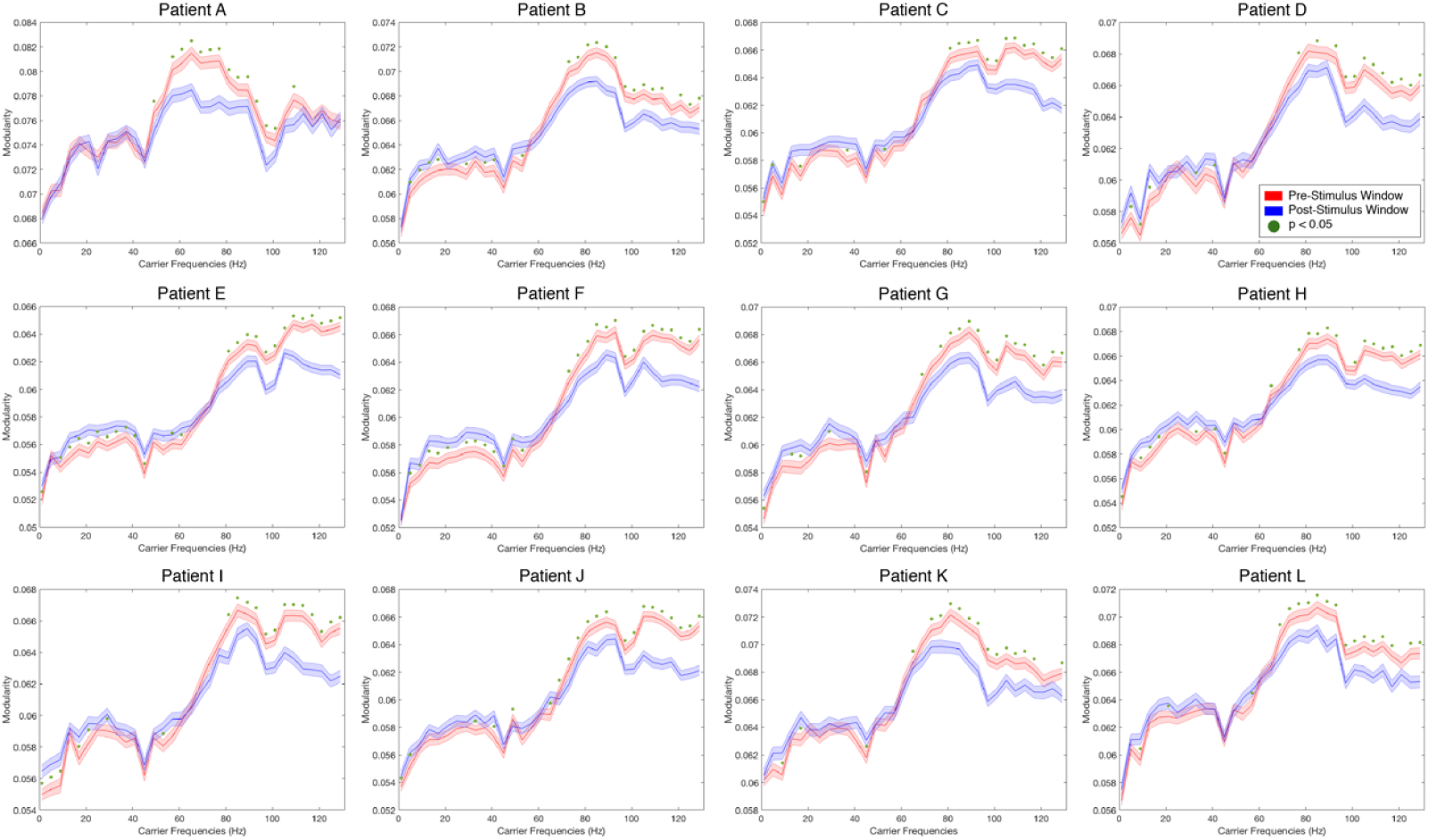
Segregation measure results for each patient. The panels show the results of segregation (measured by the modularity) during pre- and post-stimulus window for the picture-naming task. For all patients there is a significant decrease of the segregation during cognitive processing and the greatest effect can be seen in the gamma range (50 to 90 Hz). The red line corresponds to pre-stimulus window, blue line corresponds to post-stimulus window, the error shaded regions reflect the standard deviation across trials and green dots indicates a statistical significance of p<0.05 (N=1000).

**Figure 6.**
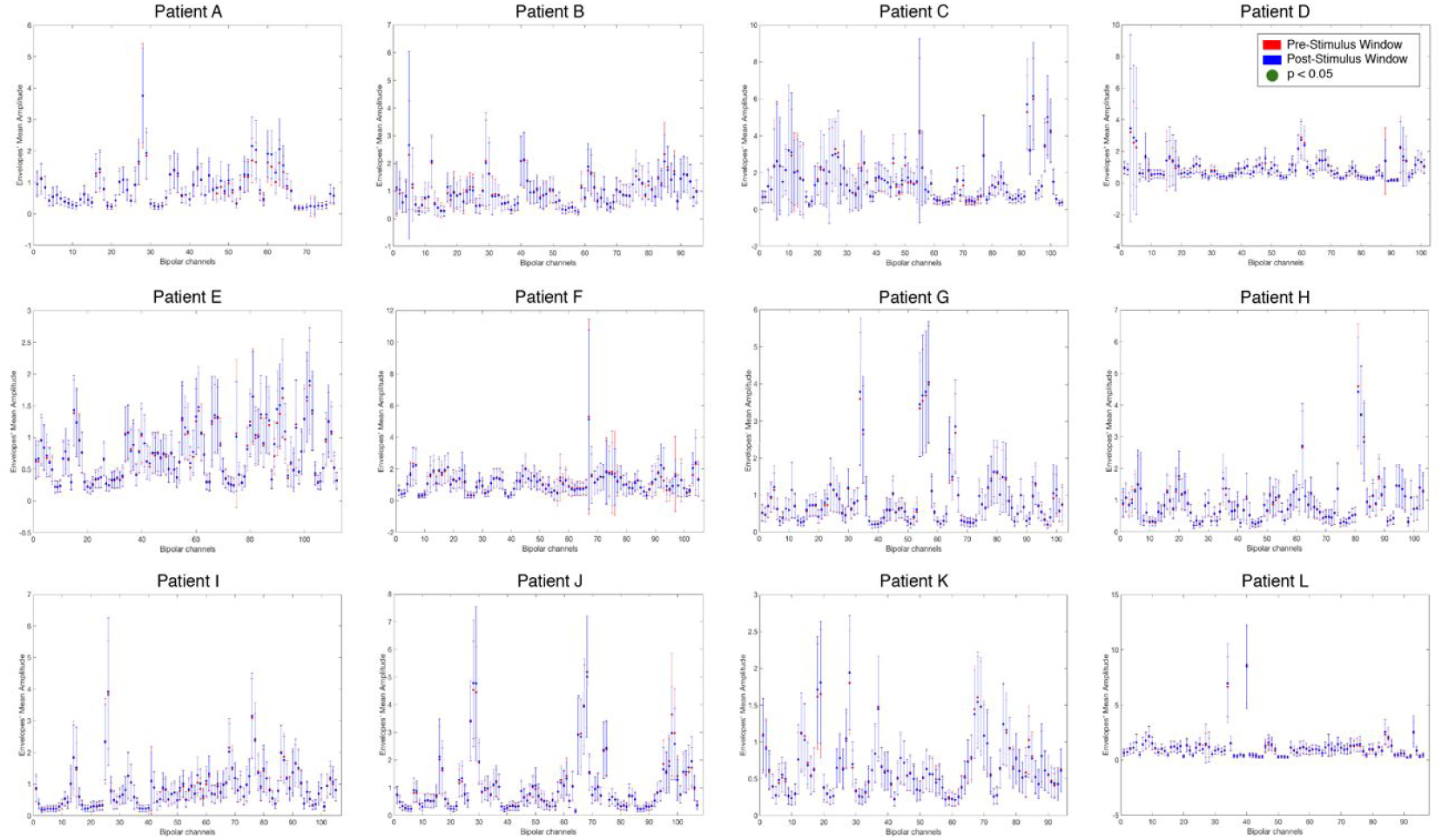
Envelopes’ Amplitude at 60 Hz across electrodes for each patient. The panel shows the results of the mean amplitude and SD at 60 Hz for each bipolar channel and both pre- and post-stimulus window for the picture-naming task. For all patients, there is no noticeable modulation across single bipolar channels between pre- and post-stimulus windows. The red line corresponds to pre-stimulus window and the blue line corresponds to post-stimulus window. The error bars reflect the standard deviation across trials. Note the strong overlap of both lines. Green dots indicate a statistical significance of p<0.05 (N=1000).

In order to uncover if the integration/segregation modulations could solely be linked to one particular task, we analysed two other cognitive tasks carried out by the patients, namely: 1) a lexical decision task; and 2) a size judgement task (see Materials and Methods). Similar to the results for the picture-naming task, we found a significant modulation of the integration post-stimulus in the post-stimulus (Figure 7A and 7D). In addition, as shown in Figure 7B and 7E we found corresponding consistent significant decreases in segregation. Consistently with the picture-naming task, the largest modulation appears in the gamma band (around 50 to 90 Hz).

**Figure 7.**
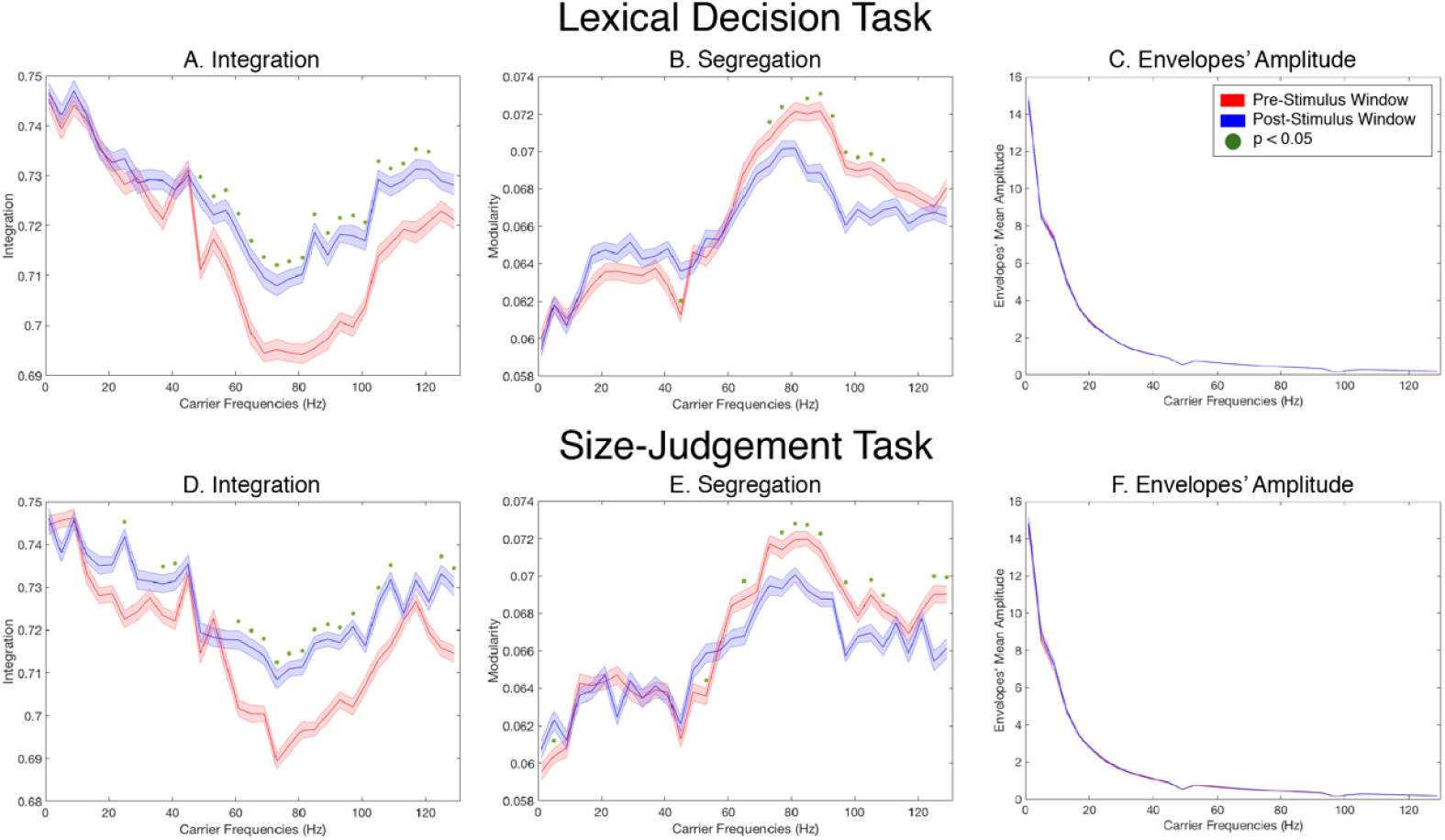
Results analysis patient K for the Lexical Decision and Size-Judgement Task. Panels A and D shows a significant increase in the integration in the post-stimulus window (red line) compared to the post-stimulus window (blue line). The greatest effect is observed in the gamma range (50 to 90 Hz). The error bars reflect the standard deviation across trials. Green dots indicate a statistical significance of p<0.05 (N=1000). Complementary to the integration measure, panels B and E shows a decrease of Segregation in the same frequency range. Panels C and F shows that there are no oscillations’ amplitude changes in any frequency induced by the stimulus (both curves are strongly overlapped). Note that the modulatory effect is almost the same for both tasks.

Further, to verify that the increases in integration and decreases in segregation measures are indeed task-driven, we contrasted the same measurements in task absence conditions. Figure 8 shows (in one participant) the lack of integration (Figure 8A) and segregation (Figure 8B) modulation when we contrasted random 50% of pre-stimulus windows trials against the other 50% (same windowing as before, i.e. 500 ms before stimulus presentation).

**Figure 8.**
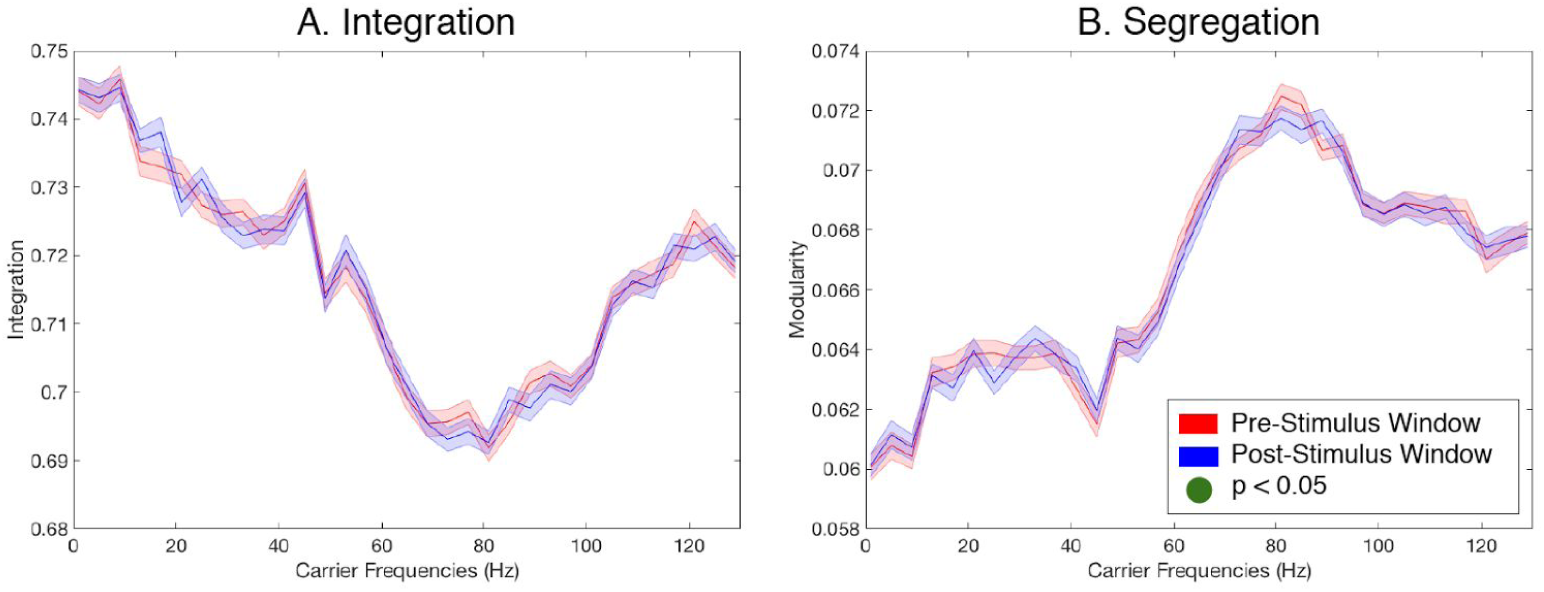
Inter pre-stimulus presentation window comparison for the picture-naming task. The panel A shows the result of the measure of Integration, and panel B shows the result of the measure of Segregation, when contrasting half of pre-stimulus trials against the other half. There is no modulation of any of the two measures before stimulus presentation. The red line corresponds to 50% of pre-stimulus windows; blue line corresponds to the other 50% of pre-stimulus window. The error bars reflect the standard deviation across trials. Green dots indicate a statistical significance of P<0.05 (N=1000).

Still, the envelope FC correlations does not necessarily reflect communication. To confirm explicitly the role of the phase coupling in the task-driven modulation of integration and segregation, we extended the analysis of those measures by using the phase-lock matrix (as a more explicit measure of communication as expressed by the CTC theory). The phase-lock matrix (see Materials and Methods) characterizes directly the global state of mutual synchronization between all pairs of network nodes. Figure 9A and 9B plots for a single patient the integration and segregation modulation during the pre- and post-stimulus windows, but now based on the corresponding phase-lock matrices (rather than the envelope FC matrices). The phase-lock matrices were computed based on the monopolar recordings. The results are consistent with the previous analysis and confirm the robustness of the findings. We also show in Figures 9C, 9D and 9E, the phase-lock matrices for a specific frequency (60 Hz, which is where we observed the greatest modulation across all patients and tasks) for pre- and post-stimulus windows and their differences respectively. The differences between both matrices (Figure 9E) demonstrate the distributed character of the modulation across many different nodes (which corresponds to each bipolar channel).

**Figure 9.**
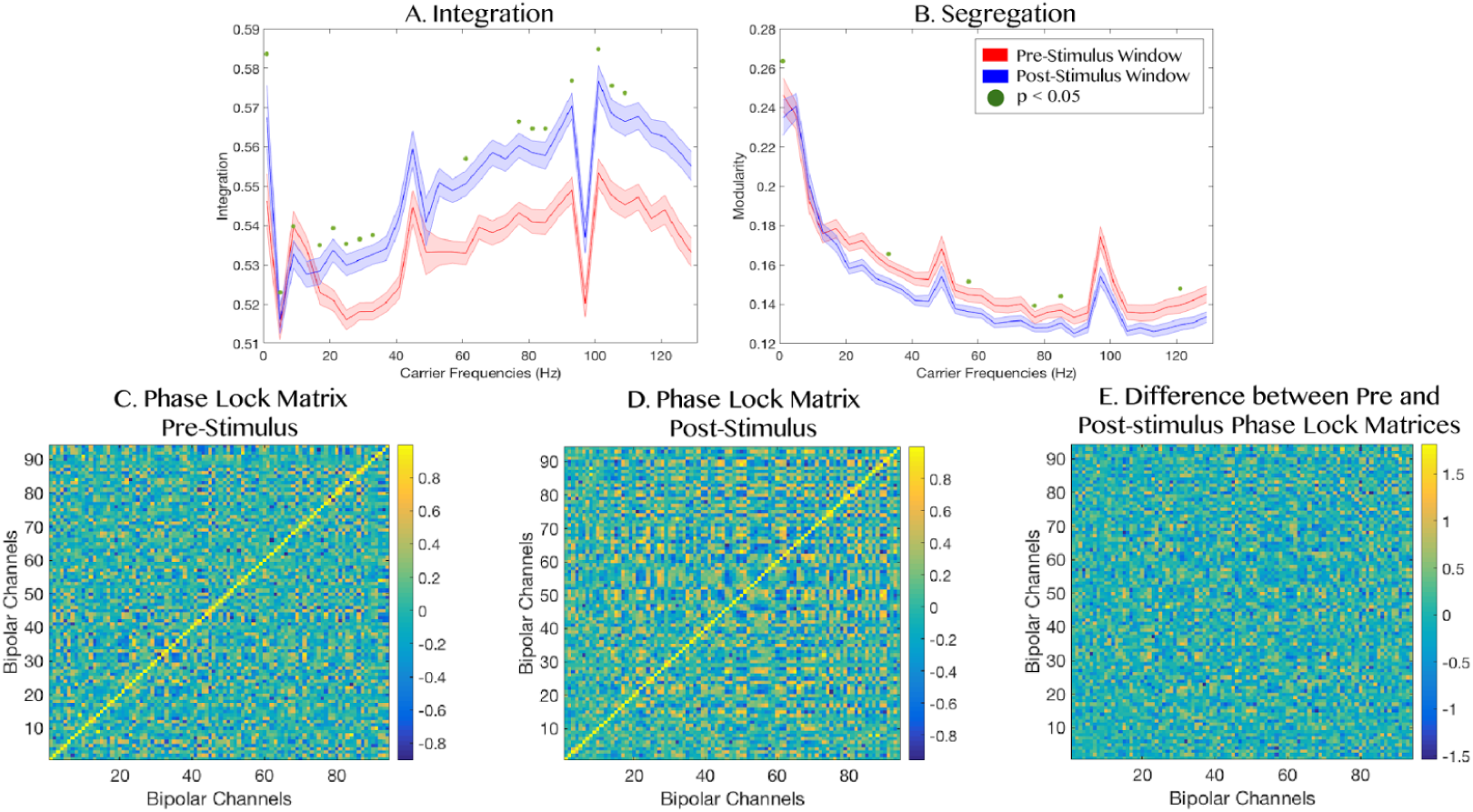
Results analysis using the Phase-lock matrix based on the monopolar montage for Patient K. Panels A and B shows the integration and segregation modulation during pre- and post-stimulus presentation, respectively, for the picture-naming task. The red line corresponds to pre-stimulus window, blue line corresponds to post-stimulus window. The error bars reflect the standard deviation across trials and green dots indicates a statistical significance of p<0.05 (N=1000). Panels C and D shows the phase-lock matrices computed for 60 Hz for the pre- and post-stimulus windows respectively. Panel E shows the difference between pre- and post-stimulus windows. The differences between both matrices (C and D, expressed on E), evidence the distributed character of the modulation across many different nodes.

Finally, we went even further to study the temporal evolution of the integration, using the phase-lock based analysis but evaluated at the millisecond level. In other words, for each single time point, we computed the phase-lock matrix (see Material and Methods) and the corresponding integration measure. Figure 10 plots the evolution of this integration measure for the whole period of time (from 1000 ms before task onset to 1500 ms after task onset). The figure shows a monotonic increasing behaviour of the integration starting after the onset of the task. This observation is relevant because it suggests that the integration is not only driven by the stimuli onset but is persistent in time. This result was found in all tasks.

**Figure 10.**
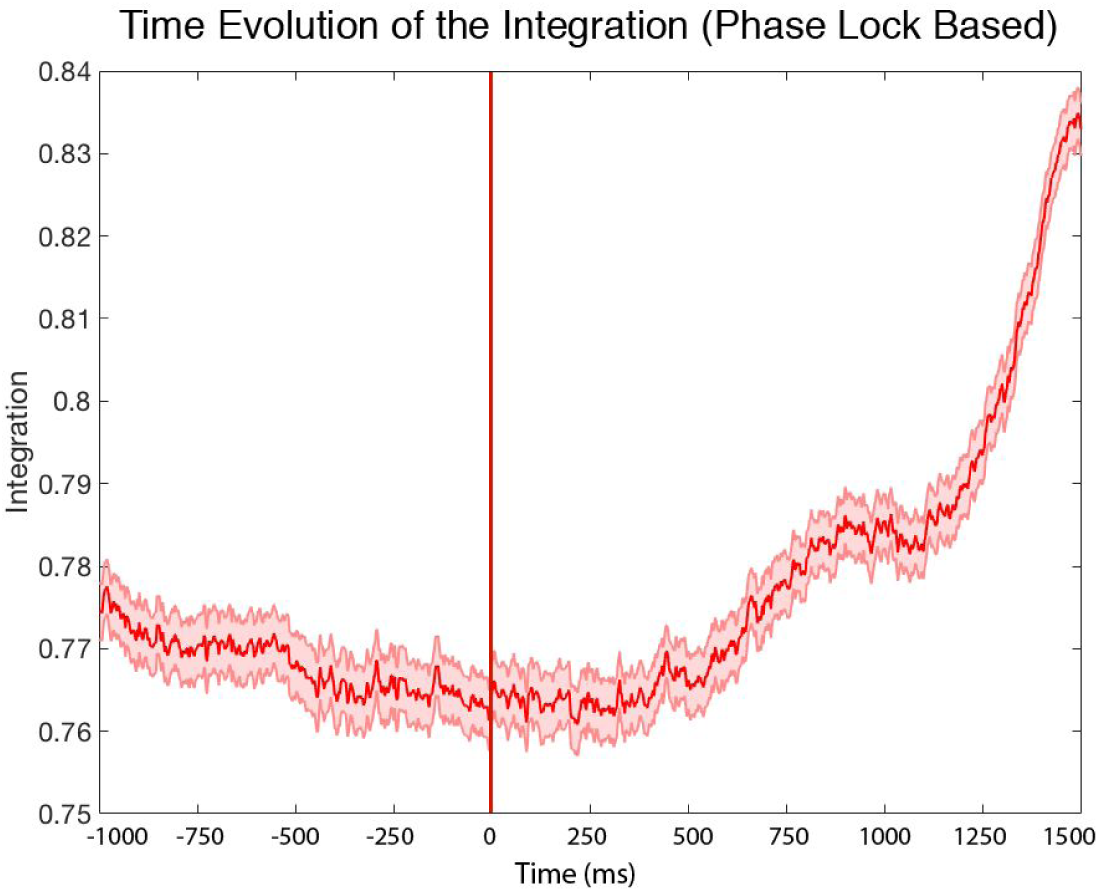
Time evolution of the Integration measure for Patient K during the picture-naming task. The figure shows the integration measure calculated for the phase-lock matrix for every single time point for the whole period of time. As seen, there is a monotonic increasing behaviour of the integration starting after the stimulus onset. The error bars reflect the standard deviation across trials.

## 4. Discussion

We used intracranial EEG (iEEG) to record human neural activity from 12 epileptic patients while they were performing three different tasks (picture-naming, lexical decision and size judgement) in order to study how cognition modulates *global* functional network measurements, namely integration and segregation. Across all patients and all tasks, we found significant increases in integration and decreases in segregation during cognitive processing (p<0.05, N=1000), especially in the gamma band (50-90 Hz). This was not driven by changes in the underlying level of oscillations given that the oscillations amplitude of pre- and post-stimulus windows were not significantly different. Instead, we found significantly higher level of synchronisation (as measured using the Kuramoto order parameter) and functional connectivity during the tasks, again particularly in the gamma band. This suggests that the cognitive modulation of the broadness of communication across the entire extended network could be due to a rearrangement of the coherence level between the nodes, which would fit the predictions of the CTC theory [38, 40] extended to the whole-brain level [37]. The CTC hypothesis posits that the mechanism through which information is transmitted is by the synchronization of distinct neuronal populations, mainly in the gamma and beta-band (30-90 Hz) [37]. Note that the original CTC theory is based on a phase synchronization mechanism defined at the neuronal level. Here, as proposed by Deco & Kringelbach [37], we extend the hypothesis to the mesoscopic/macroscopic level by analysing the phases and correlations of the envelopes of the Band Limited Power.

Our results support the view of the brain as a complex system organized into large-scale networks. In order to support cognitive functions, the networks need to flexibly adjust their functional connectivity in order to integrate relevant information to support goal-directed behaviour. Graph and information theoretical approaches have helped to characterize the global network connectivity in terms of *segregation* and *integration* [22, 53, 54]. Here, segregation refers to the relative statistical independence of subsets of neurons or brain regions to compute information [53, 55], while integration is a complementary concept quantifying the level of connectivity across the whole-brain [22, 55]. In particular, we used a measure of integration based on the largest component present in the whole-brain connectivity matrix [22], while for the segregation we used the concept of modularity [56].

Our findings are in line with previous neuroimaging studies finding increases of the integration across brain networks as evidenced by MEG during working memory processing [25, 36], or by MRI during emotional and motivational stimulus processing [31] and fMRI for selective attention [57].

We found that there was a decrease in segregation in the gamma band with task, yet an increase in segregation in the sub-gamma band. The decrease of the segregation in the gamma regime is expected and consistent with the increase of integration in the same regime which is putatively linked to increased stimuli/cognitive processing [40]. Along the same lines of argument, the increase of segregation in the sub-gamma bands could perhaps be associated with a disengagement of large slow resting networks. It would be of considerable interest to further investigate.

Interestingly, the increase of the integration level was persistent after the onset of the task suggesting that the integration is putatively not only driven by the stimuli but related to the cognitive processing.

It is important to remark the fact that the implantation scheme of the electrodes is different for each patient. This works to our advantage, since the main aim was to show whether there is an increase of integration (i.e. increases in the level of connectivity across the whole network) across the whole-brain, and thus independent of the positions of individual electrodes. Further strengthening the results, we analysed three different cognitive tasks, which demonstrated that the main modulatory effect on the integration/segregation and the validity of the CTC theory was independent of the particular cognitive processing. Thus, because of our global perspective, unlike many previous studies, we did not investigate the role of single channels in the different cognitive processes. Nevertheless, we are convinced that localising these electrodes in the human brain could lead to the discovery of interesting differences at the local spatial level and that this could be a very interesting goal for future investigations.

Equally, it would be interesting to examine in future investigations how the integration and segregation measures might be differently modulated under different cognitive load and as a function of the task were performed correctly or erroneously. An important future goal would be to go beyond global measures to determine specifically which brain areas or local networks show the highest degree of task-driven effective connectivity and therefore are mostly involved in information processing. In this context, future studies could greatly benefit from diffusion tensor imaging in twofold ways: to visualize and describe the precise connectivity between the electrodes in the brain, as well as the basis of whole-brain models considering the connectivity and continuity of neural pathways in the patients.

There are certain methodological limitations due to the fact that iEEG in humans is always obtained from epileptic patients who, besides having epileptogenic neural activity, may have also differences in the structural connectivity. On the other hand, the lack of whole-brain coverage with this technique means that we have to care when making assumptions of global network connectivity because of the restriction of the spatial coverage that can be simultaneously sampled.

## 5. Conclusions

We show that cognitive processing drives a global increase of integration and decrease of segregation, especially in the gamma band (50-90 Hz), which could reflect broadcasting of communication. The results add to the emerging interesting literature on integration and segregation in the human brain (see [58] for a carefully review). More importantly, we demonstrate here for the first time that these modulations in the level of functional network topology were not caused by changes in the level of the underlying oscillations but caused by a rearrangement of the mutual synchronisation between the different nodes, according to the “Communication Through Coherence” Theory.

## 6. Acknowledgements

The authors would like to thank all patients for their participation and the IMIM-Hospital del Mar Epilepsy Unit staff for their technical assistance in collecting the data. This work was supported by Gustavo Deco’s ERC Advanced Grant: DYSTRUCTURE (n. 295129), from the European Union’s Horizon 2020 research and innovation programme under grant agreement No. 720270 and by the Spanish Research Project PSI2013-42091-P. M.L.K. is supported by the ERC Consolidator Grant CAREGIVING (n. 615539) and the Center for Music in the Brain, funded by the Danish National Research Foundation (DNRF117).

